# Widespread Receptor Driven Modulation in Peripheral Olfactory Coding

**DOI:** 10.1101/760330

**Authors:** Lu Xu, Wenze Li, Venkatakaushik Voleti, Elizabeth M. C. Hillman, Stuart Firestein

## Abstract

We utilized swept confocally aligned planar excitation (SCAPE) microscopy to measure odor-driven activity simultaneously in many (>10,000) olfactory sensory neurons distributed over large areas of intact mouse olfactory epithelium. This approach allowed us to investigate the responses to mixtures or blends of odors and their components, a more realistic stimulus than monomolecular odors. In up to 38% of responding cells, responses to a mixture of odors were different - absent, smaller or larger - than what would be expected from the sum of the individual components. Further investigation revealed instances of both antagonism and allosteric enhancement in the primary olfactory sensory neurons. All 10 of the odor compounds tested were found to act as both agonists and antagonists at different receptors. We present a hypothetical scheme for how modulation at the peripheral receptors increases the capability of the olfactory system to recognize patterns of complex odor mixtures. The widespread modulation of primary sensory receptors argues against a simple combinatorial code and should motivate a search for alternative coding strategies.

---

With the seminal discovery of a very large number of olfactory receptors^1–5^ it appeared that a combinatorial, additive code, not unlike color vision, would be the most likely model for encoding olfactory sensory inputs. For monomolecular stimuli it is indeed possible to identify subsets of receptors that respond to particular odors or groups of related odors, and which could serve to unambiguously discriminate one odor from another^6–9^. However, under realistic conditions the olfactory system must handle a far more complex input, consisting primarily of blends of odors that may number in the many hundreds. Among those varied molecules there may be some that act as modulators as well as simple agonists. Suppression in blends has been known from psychophysical experiments for decades^10–14^, but the neural locus of those effects remains largely unknown (unlike vision where inhibition plays a large role, but only in higher circuitry). In the current study we utilized a new technology, Swept Confocally Aligned Planar Excitation (SCAPE) microscopy^15–17^. SCAPE enabled high-throughput, high-speed 3-dimensional, simultaneous monitoring of the intracellular calcium responses of large numbers of individual intact olfactory sensory neurons (OSNs) expressing GCaMP6f, imaged in-situ within a uniquely designed mouse hemi-head perfused preparation. Observations of responses of the same cells to a sequence of pure odors and then to mixtures of the same odors revealed a significant number of cells whose activity was modulated by one or more components of the mixture. This modulation could take the form of suppression or enhancement, consistent with both antagonistic and allosteric effects at the primary receptors. These results call into question what is likely an overly simple model of a linear combinatorial code for discrimination of odors or odor mixtures. An additional and unexpected finding to come from these data is the occurrence of small molecule allosteric modulation of these Class A GPCRs.

## RESULTS

Previously several laboratories have shown the possibility of antagonistic inhibition at olfactory receptors^8,18–23^. Given that they are members of the Class A family of GPCRS, this is not entirely surprising. However, the role of inhibition, or other modulatory effects on the primary receptors has not been extensively examined in an intact preparation. We used a hemi-head, intact mouse preparation to image the activity of olfactory sensory neurons during perfusion of individual and combinations (blends) of specific odors. These mice expressed the genetically encoded Ca^2+^ indicator GCaMP6f under the mature OSN-specific OMP promoter (OMP-GCaMP6f). The physical layout of the preparation can be seen in Fig. 1A and the details are provided in Methods. This preparation enabled a variety of odor stimuli to be perfused over the intact tissue while tracking the simultaneous responses of thousands of olfactory neurons layered within the epithelium of the curved turbinates (Fig. 1B, C). Extended Data Movie 1 shows an example of dynamic neural responses to an odor stimulus. Mature OSNs express only one allele of an odor receptor gene ^24–27^, and therefore the response of a single neuron should represent the response of its receptor. With this new technique we were able to detect activity evoked by the application of the olfactory cell adenylyl cyclase activator, forskolin, in nearly 10,000 OSNs per mouse.

**Figure 1.**
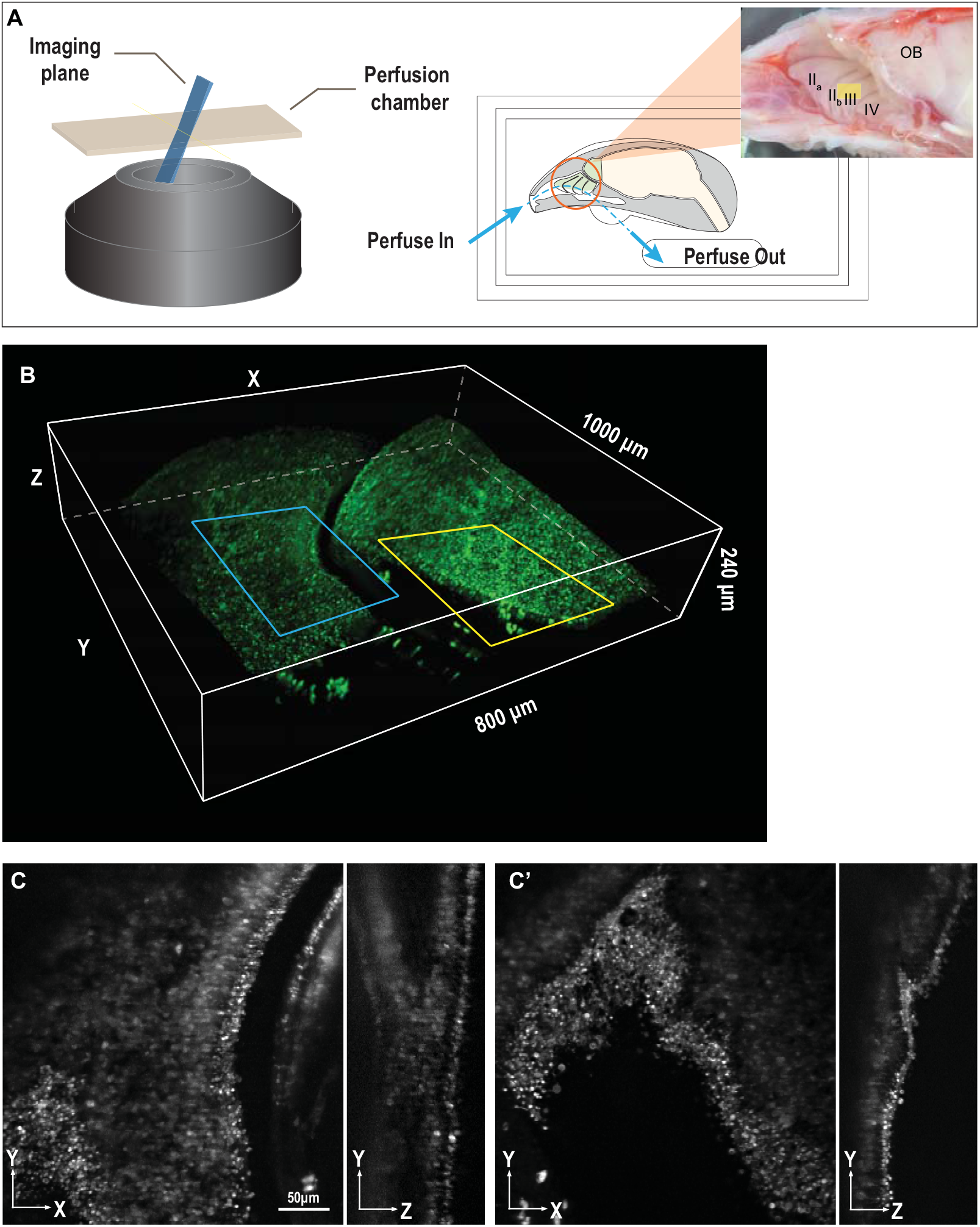
Imaging of intact olfactory epithelium using SCAPE microscopy. **A.** Schematic of intact olfactory epithelium imaging platform for SCAPE microscopy. The SCAPE microscope objective is configured in an inverted layout. A custom-designed glass bottom perfusion chamber was placed above the objective with water immersion. The right half of mouse head was mounted in the perfusion chamber, with olfactory turbinates exposed. The perfusion chamber was designed to control the perfusion flow through the nasal cavity with the inlet at the nostril and the outlet at the throat (blue arrows). The field of view typically covered the ventral half of either turbinate IIb or III, and some of the neighboring turbinates (yellow rectangle). **B**. 3D volumetric rendering of SCAPE data acquired on the olfactory epithelium showing a 1000μm×800μm×240μm field of view. The same volume could be imaged for more than an hour at 2-5 volumes per second (VPS) in the perfusion chamber during sequential odor delivery. **C, C’**. Top-down and side view of the highlighted regions (blue in C and yellow in C’). Maximum intensity projection of 10μm sub-stack in Z and 5μm sub-stack in X were shown here. Scale bar = 50μm. Note that individual olfactory sensory neurons (OSNs) can be resolved.

### Odor blends

To test OSN responses to odor blends, we designed two odor sets: odor set 1 (Fig. 2A) contained Acetophenone, Benzyl acetate and Citral, all at 100μM. These odors are common ingredients in perfumery but possess different nuances: Acetophenone, the simplest aromatic ketone, is often described as almond or mimosa; Benzyl acetate is described as possessing a floral and jasmine scent, and Citral is described as citrus. Thus the three components of the mixture are chemically and perceptually distinct (at least to humans). Additionally the three odors were chosen because they can activate a large number of cells for analyses. Odor set 2 (Fig. 4A) is a specific formula well known to perfumers as a “woody accord” and contains 148μM Dorisyl, 48μM Dartanol and 127μM Isoraldeine. It was chosen both because it is a common blend with a singular perception (again to humans) and it is chemically quite different from mixture 1. Stimuli were pseudo-randomly presented as the total mixture, each component singly and in the 3 possible binary pairs.

**Figure 2.**
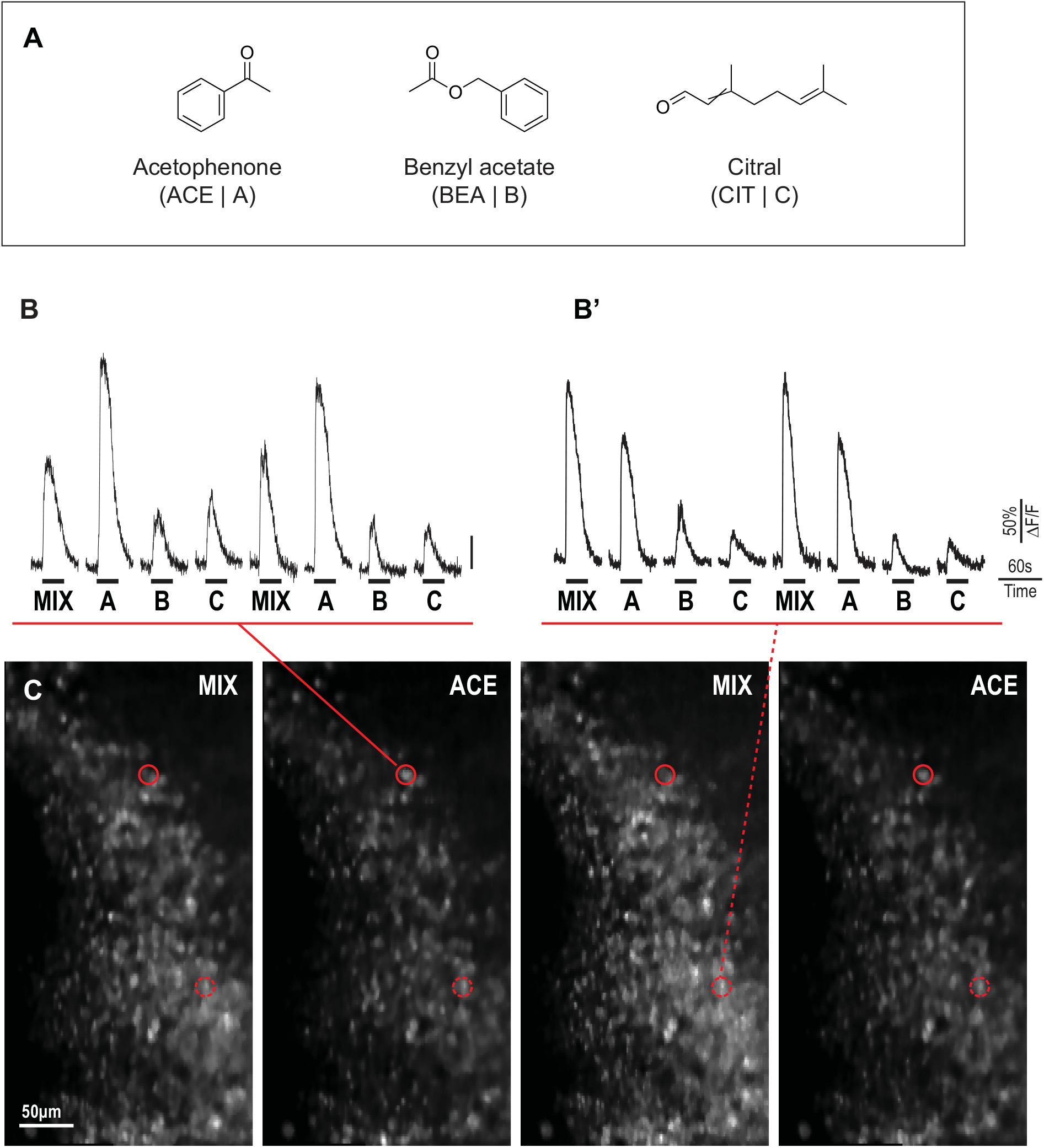
Representative GCaMP time courses of individual OSNs extracted from raw SCAPE image. **A.** Chemical structures of the three odorants used as odor stimuli (odor set 1). **B, B’.** Two sample GCaMP time courses of individual OSNs. Time course was extracted directly from the raw volumetric time series imaged by SCAPE microscopy. A circular ROI centered at each individual OSN with 7.7μm diameter was used to extract the GCaMP activity. A 30s-long stimulus was delivered in each trial, with a 2.5min interval (not shown) between stimulus applications. **C.** A subset of Acetophenone responsive OSNs are inhibited by components of the mixture. From this maximum intensity projection of a 7.7μm-depth sub-stack cropped from a 1000μm×500μm×200μm volume imaged by SCAPE microscopy two OSNs are circled in red. Both of these neurons responded to Acetophenone. One of them however was inhibited by the mixture of which Acetophenone was one component (solid line) while the other was unaffected by the mixture (dashed line). The raw time courses of these neurons are shown in figure B and B’. Each volume was recorded at 2s after the odor stimulus arrived in each trial. Scale bar = 50μm.

### Single neuron activity extraction from SCAPE imaging

Dynamic 3D SCAPE images of epithelium responses were analyzed after performing motion correction over the whole trial. 11 to 18 trials (75s each) of volumetric recording were divided into several 7.7μm thick depth sub-stacks and concatenated together for further data analysis. Time-courses from each neuron were then extracted using 2D constrained nonnegative matrix factorization (CNMF) from the sum intensity projection of each sub-stack to obtain single cell calcium time courses and cell locations ^28^.

### Responses to odor set 1

Recording the number of cells responding to the mixture and then to each of the individual odors showed that the sum of the cells responding to each odor was greater than the number of cells responding to the mixture. One simple explanation for this is that some cells are activated by more than one of the odors and therefore are counted twice in the individual cell census. However, there were clear instances of cells that responded to one of the odors but gave either a reduced or no response when presented with the mixture containing the same odors at the same concentrations. Fig. 2C shows an example of one such cell (solid red circle) that showed a high response to Acetophenone alone and a significantly reduced response to the Acetophenone containing mixture. The trace above the image (Fig. 2B) shows the responses of the cell. In these cases, which were numerous (see later), it appeared that the most reasonable explanation was that one of the other components in the mixture was acting as an antagonist at certain receptors.

To determine if this was indeed the case, and to obtain a more comprehensive neuronal response profile, we sampled a large number of cells with the mixture, with each component of the mixture individually and with each of the three pairwise combinations of the three odorants. Fig. 3A shows data from more than 10,000 cells where peak responses were normalized and plotted in a heat map format. Each row represents the response of a single cell and each column represents an odor stimulus condition (**Mix**-**A**-**B**-**C**-**Mix**-**AB**-**AC**-**BC**-**Mix**, where **A/B/C** stands for Acetophenone, Benzyl acetate and Citral). The data are sorted into subgroups based on response patterns using k-means clustering. Eight major subgroups can be observed from top to bottom: cells not (or barely) activated by any of the stimuli, cells dominantly activated by one of the three individual odors, cells dominantly activated by a binary pair of two of the three individual odors, and cells activated by all three individual odors. Representative time courses of each subgroup can be seen in Extended Data Fig. 1.

**Figure 3.**
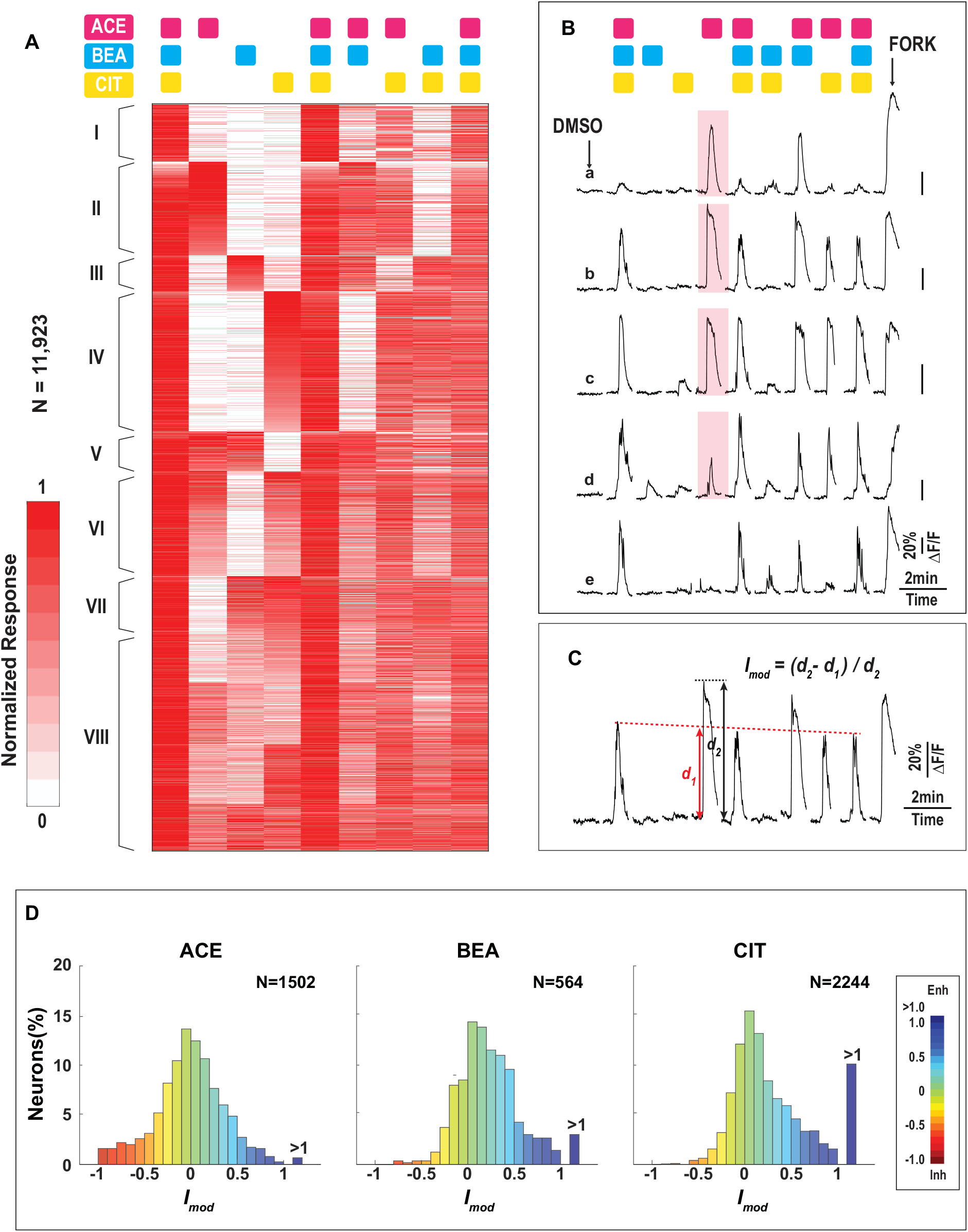
Response profile of odor set 1. **A.** Heatmap of normalized peak responses (N = 11,923, 5 mice) to odor set 1. Odor stimuli (columns) were given in a pseudo-stochastic manner for each mouse and re-aligned for this presentation. OSNs (rows) were clustered into 8 subgroups (I-VIII) based on k-means clustering, as detailed in the text. Odor stimuli combinations are denoted by the colored squares at the top. **B**. Representative time courses of individual OSNs showing inhibition/suppression (a,b; Acetophenone primarily by Citral), no effect (c) and enhancement (d,e; Acetophenone by either Benzyl acetate alone or both Benzyl acetate and Citral) respectively. These traces are all from cells in Groups I and II of the Heatmap (see Fig. 3A). Additional representative traces for all 8 groups are shown in Extended Data Fig. 1 which also provides the basis for the groupings. **C**. Quantification of the modulation effects. For neurons that were most responsive to one odorant (Acetophenone in this case), a modulation index (*I_mod_*) was calculated as (*d_2_*-*d_1_*)/*d_2_*, where *d_2_* was the response magnitude to the single odorant, and *d_1_* was the response magnitude to the mixture linearly corrected based on responses to the flanking **Mix** stimuli (see Methods). **D**. Representations of the modulation effects on Acetophenone /Benzyl acetate /Citral dominant neurons in the left, middle and right histograms respectively. These are subgroups II, III, IV in Fig. 3A. Red indicates that cell responses to the individual odors were inhibited by the mixture and blue indicates enhancement by the mixture. 18% / 3% / 3% neurons in each subgroup showed >30% suppression; 19% / 38% / 38% neurons showed >30% enhancement, respectively.

Among the subgroups, cells dominantly activated by one of the three individual odors (Fig. 3A, subgroups II-IV) were of particular interest because these cells provided a straightforward comparison between responses to that individual odor and to the mixture. For example, in the subgroup (II) that dominantly responded to Acetophenone (**A**) most cells responded at the same magnitude to **A**, **AB**, **AC** and **Mix**, indicating that Benzyl-acetate (**B**) and Citral (**C**) neither co-activated these cells nor modified their responses to Acetophenone (Fig. 3B, cell c). However, there were important exceptions, as some cells responding to Acetophenone were suppressed or even completely inhibited (cell a and b) by the mixture, or as in the case shown, by the Citral component in particular, while responses in other cells were enhanced (cell d). Similarly, instances of suppression/enhancement were also observed in the Benzyl-acetate and Citral dominant groups (III and IV). To quantify these effects, a modulation index (*I_mod_*) was calculated, where negative values indicate suppression and positive values indicate enhancement (Fig. 3C). The distribution of modulation effects for each odorant are plotted in Fig. 3D. These effects center around 0, no effect, as would be expected. Nonetheless there is a significant degree of modulation, including both suppression and enhancement, even between only these three odors. (The percentage of enhancement may be slightly over-estimated because some single odor-dominant OSNs also responded weakly to the other two odors. However, in many cases (29%, 31% and 54% in groups II, III, IV), the response to the **Mix** is still higher than the numeric sum of the responses to the individual odorants.) In a control experiment in which only the **Mix** was delivered repetitively the *I_mod_* had a variance of only 9% (Extended Data Fig. 2, see Methods).

Interestingly, the top most subgroup (Fig. 3A, subgroup I) shows cells that did not have a significant response to any of the individual odors but responded strongly to the mixture (Fig. 3B, cell e). There are two explanations for this behavior. The simplest is that the receptors in these cells responded to each of the three odors, but only at high concentrations, and the **Mix** represented a 3-fold higher total concentration than the highest concentration tested individually (300μM vs. 100μM). Thus, these could simply have been relatively insensitive cells that were activated by the higher total concentration of odors in the **Mix**.

While this seems a straightforward explanation it does require that this group of cells expressed receptors that would have had at least some affinity for all three odors in the **Mix** and that the three odors act independently at the receptor. Given the structural differences between the odors this seems unlikely and would itself be remarkable. We believe that a reasonable alternative explanation is that one or more of the odors in the **Mix** was indeed a weak agonist and that one or more of the odors provided an allosteric enhancement. We will provide additional evidence for this possibility below.

### Responses to odor set 2

The experimental design here is similar to that of odor set 1 with the major exception being that the components of the mix are not all at the same concentration. This blend was designed by perfumers to produce a woody perception and both the components and the concentrations have been worked out to maximize that effect. While perfumers apparently arrived at this formula by experience and tacit knowledge, we sought to determine if suppression and enhancement might play a role in their choices. In this blend we noted that a significant number of Dorisyl-dominant neurons were suppressed (Fig. 4B, black box). OSN responses to the binary mixtures further indicated that Isoraldeine (asterisk) was the major inhibitor rather than Dartanol (triangle). The modulation profile is summarized in Fig. 4C.

**Figure 4.**
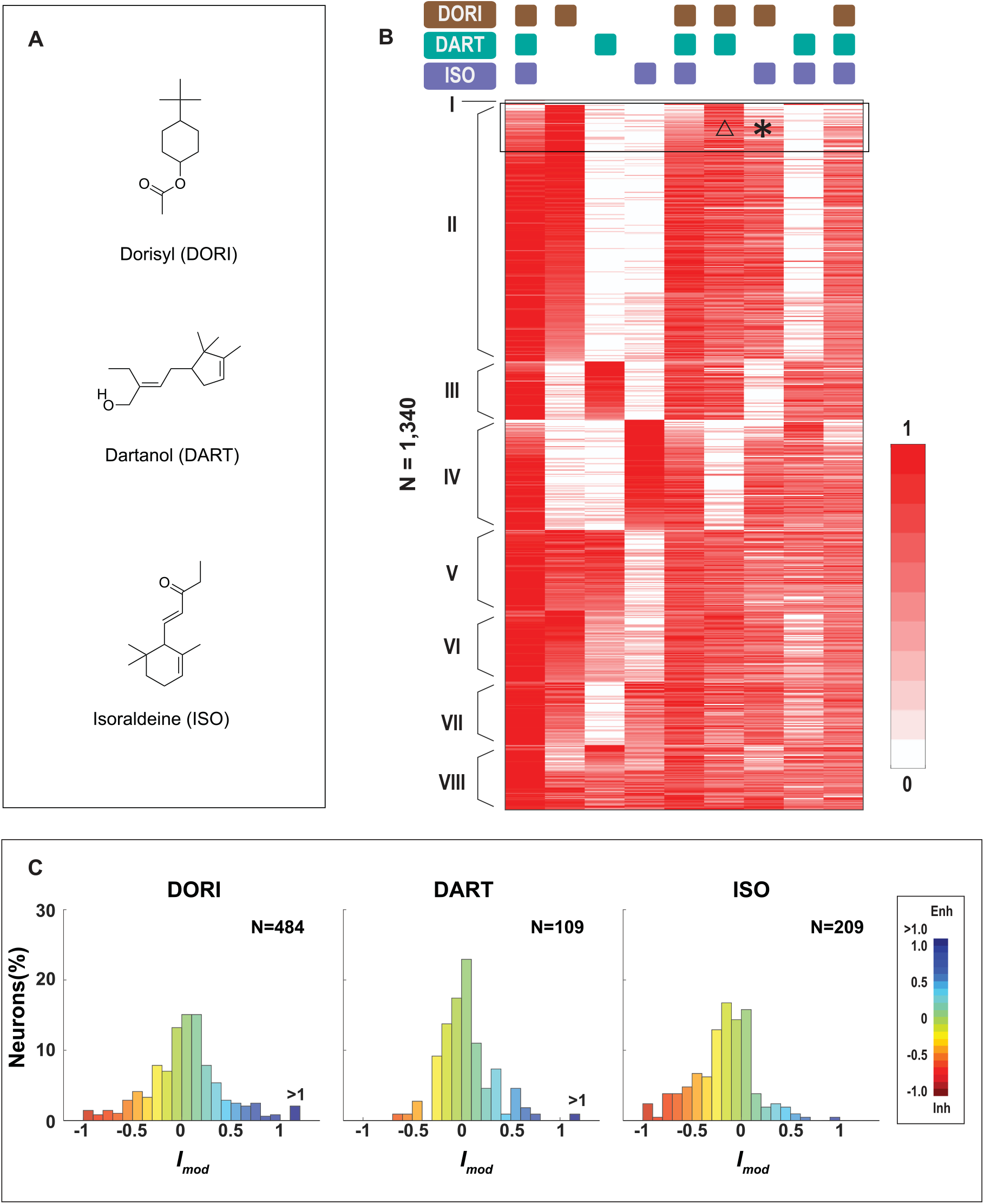
Response profile of odor set 2. **A**. Chemical structures of components in odor set 2 (the woody accord). **B**. Heatmap of normalized peak responses (N = 1,340, 3 mice) to the woody accord. Odor stimuli (columns) were given in a pseudo-stochastic manner and re-aligned for presentation clarity. The 8 groups were determined in the same manner as for Odor mix 1. **C**. Histograms showing the distribution of modulation effects. Dorisyl/Isoraldeine/Dartanol-dominant neurons were analyzed and plotted in left, middle and right figures, respectively. Red indicates cell responses to individual odors were inhibited by the mixture and blue indicates enhancement. 15% / 5% / 28% neurons in each subgroup showed >30% suppression; 19% / 17% / 6% neurons showed >30% enhancement, respectively.

### Dose response analysis shows that inhibition is due to partial or competitive antagonism

To gain a deeper understanding of the mechanism of the observed modulation effects we undertook a series of dose-response experiments using the responses of single cells. With SCAPE we were able to sample a large number of cells for these experiments. Acetophenone was selected as the agonist and Citral as the modulator – that is, we selected cells that were activated by Acetophenone and showed little or no response to Citral.

7,030 OSNs were activated by either component in this binary odor pair (Extended Data Fig. 3A). After k-means clustering and data sorting, 168 OSNs that responded to Acetophenone were found to be suppressed (by >30%) or completely inhibited by 100μM Citral (Fig. 5A). Single-neuron responses showed that neuronal responses were more likely to be inhibited at lower Acetophenone concentrations; as Acetophenone concentration increased, the suppression effect could be overcome (Fig. 5B). Normalized responses of all 168 OSNs were plotted against Acetophenone concentration. The suppression effect of Citral was dependent on Acetophenone concentration, suggesting competitive antagonism (Fig. 5C). In a few cases (15/168) Citral alone could also activate an OSN but would nevertheless suppress the activity of Acetophenone (Extended Data Fig. 3B). These cases fit the standard model of partial agonism, which we also observed regularly. Similarly, dose-dependent inhibition was also observed in the Dorisyl-Isoraldeine odor pair, where Dorisyl was the agonist and Isoraldeine was the antagonist (Extended Data Fig. 4).

**Figure 5.**
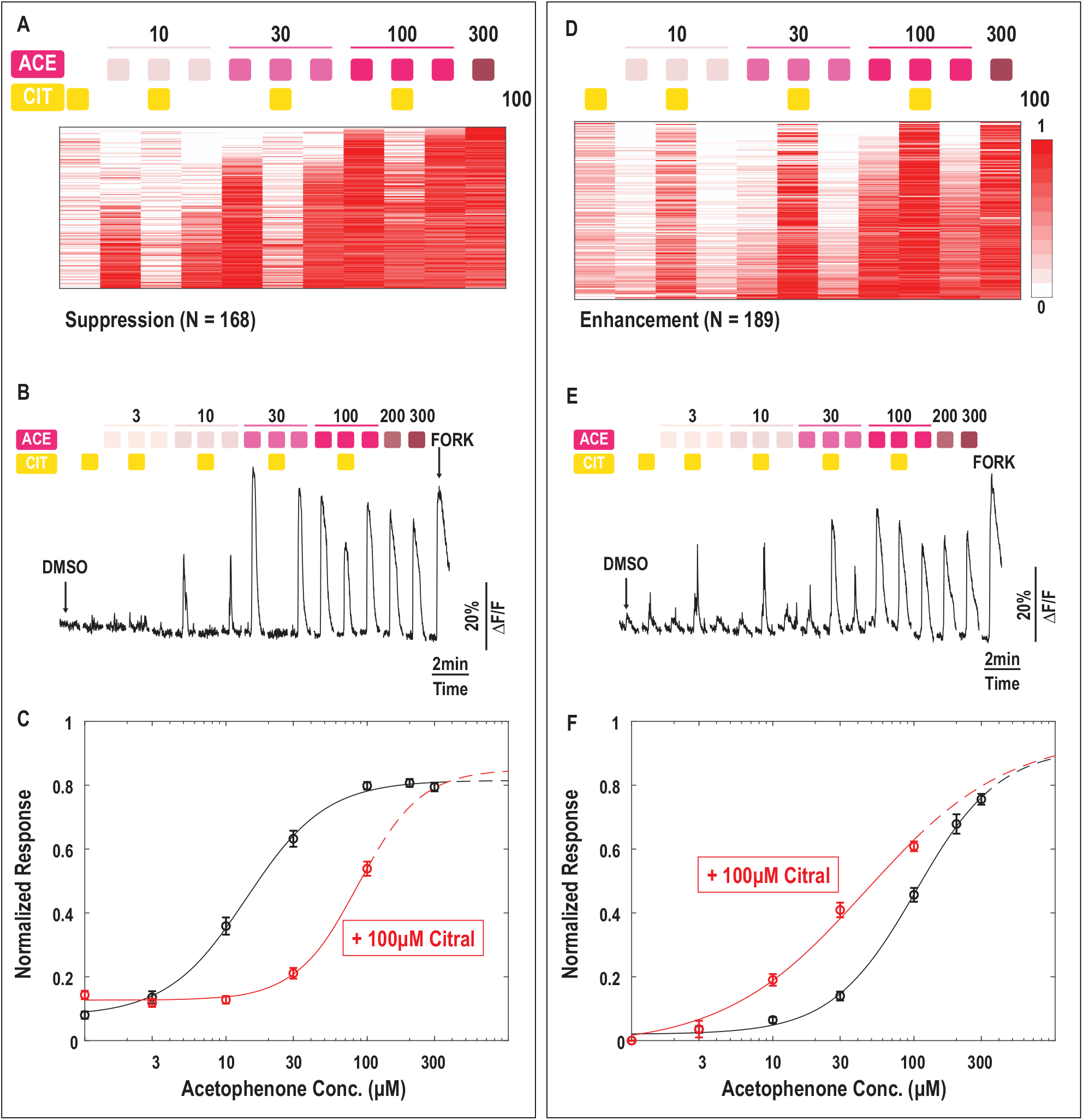
Dose-dependent suppression/ enhancement of Acetophenone by 100μM Citral. **A.** Normalized response heatmap of Acetophenone-activated neurons suppressed by Citral (N = 168). Neurons were stimulated by an increasing concentration of Acetophenone (10-300μM) in the presence or absence of 100μM Citral. OSNs that showed >30% suppression were considered as suppressed. Odor stimuli were denoted by colored squares at the top. **B**. Representative time course of an individual OSN suppressed by 100μM Citral. **C**. The effect of 100μM Citral as an antagonist. OSNs suppressed by 100μM Citral (see Fig. 5A) were selected for this analysis. Dose-dependent responses (mean±S.E.M.) were plotted and fitted with the Hill Equation. Responses to Acetophenone alone were plotted in black; responses to Acetophenone + Citral were plotted in red. Acetophenone alone: Hill coefficient = 1.55, EC_50_ = 13.99μM; Acetophenone + Citral: Hill coefficient = 1.95, EC_50_ = 86.53μM. **D**. Normalized response heatmap of Acetophenone-activated neurons enhanced by Citral (N = 189). OSNs that showed >30% enhancement were considered as enhanced. **E**. Representative time course of an individual OSN showing enhancement. **F**. The effect of 100μM Citral as an enhancer. To avoid bias, the baseline responses to 100μM Citral /DMSO were linearly subtracted from the dose-dependent responses (mean±S.E.M.) to Acetophenone + Citral (red) and Acetophenone alone (black), respectively. Results were fitted with Hill Equation. Acetophenone alone: Hill coefficient = 1.46, EC_50_ = 102.83μM; Acetophenone + Citral: Hill coefficient = 0.87, EC_50_ = 45.42μM.

### Enhanced Responses

While competitive antagonism and partial agonism are not surprising mechanisms to find in themselves among GPCRs, enhancement of responses is harder to explain. Nonetheless we recorded 189 OSNS whose responses to Acetophenone were significantly (by >30%) enhanced by 100μM Citral (Fig. 5D). In most of the cases we observed a low concentration of Acetophenone was insufficient to activate OSNs, unless mixed with 100μM Citral (which produced only a small response on its own), while high concentrations of Acetophenone (200 or 300μM) were sufficient to activate these OSNs alone (Fig. 5E). A dose-response plot of these OSNs (Fig. 5F) shows a clear shift to the left (enhancement) caused by the presence of Citral. In some cases (n=5 in this data set), even 300μM Acetophenone could not elicit a response, whereas the mixture of lower concentrations of Acetophenone and 100μM Citral could activate the neuron (Extended Data Fig. 3C). We suggest, from these data, that the mechanism for this enhancement is most likely to be an allosteric effect of Citral on this subset of receptors. The combined concentration of 100μM Citral and 100μM Acetophenone (total equals to 200μM) gives a larger response than 200μM or even 300μM Acetophenone alone. Therefore the enhanced effect cannot be due only to increased ligand concentration. Although small molecule allosteric modulation of Class A GPCRs has been only rarely observed^29–33^, the wide diversity of the OSN receptor family appears to have extended the occurrence of this mechanism in GPCRs of this type. We are unable from these experiments to determine the allosteric site, but we note that the odor molecules are relatively hydrophobic and from a chemical point of view, could easily insert themselves in the lipid membrane and bind to sites within the transmembrane regions of the receptor.

### How widespread is receptor modulation? Screening of additional Acetophenone modulators

While we have used a simple blend of a just a few odors and further concentrated on one or two of them as modulators, there is nothing inherently special about any of these molecules. Indeed we suspected that virtually all odors can act as both agonists and modulators at various receptors, depending on what other molecules may be present. This response modulation could occur between any pair of odors that happen to activate/antagonize/enhance the same receptor. If this is correct, Citral should not be the only antagonist of Acetophenone, a prediction that can be tested empirically.

To test this hypothesis, we selected four different odorants (Dartanol (DART), Isoraldeine (ISO), γ-terpinene (gTER) and Isoamyl acetate (IAA); see Fig. 6A) and paired them with Acetophenone (ACE). We recorded 6,178 cells that were activated by at least one of the odors (Extended Data Fig. 5); among which 1,309 were activated by Acetophenone. After k-means clustering and cell sorting, 85 cells showed at least a 30% response suppression by one or more of the other compounds, Isoraldeine being the most common (Fig. 6B). Interestingly, the responses to the four compounds were varied, indicating that multiple types of receptors were likely involved (Fig. 6C, cell a and b). In rarer cases, cells were inhibited by more than one odorant (cell c), and sometimes were even suppressed by all four odorants equivalently (cell d).

**Figure 6.**
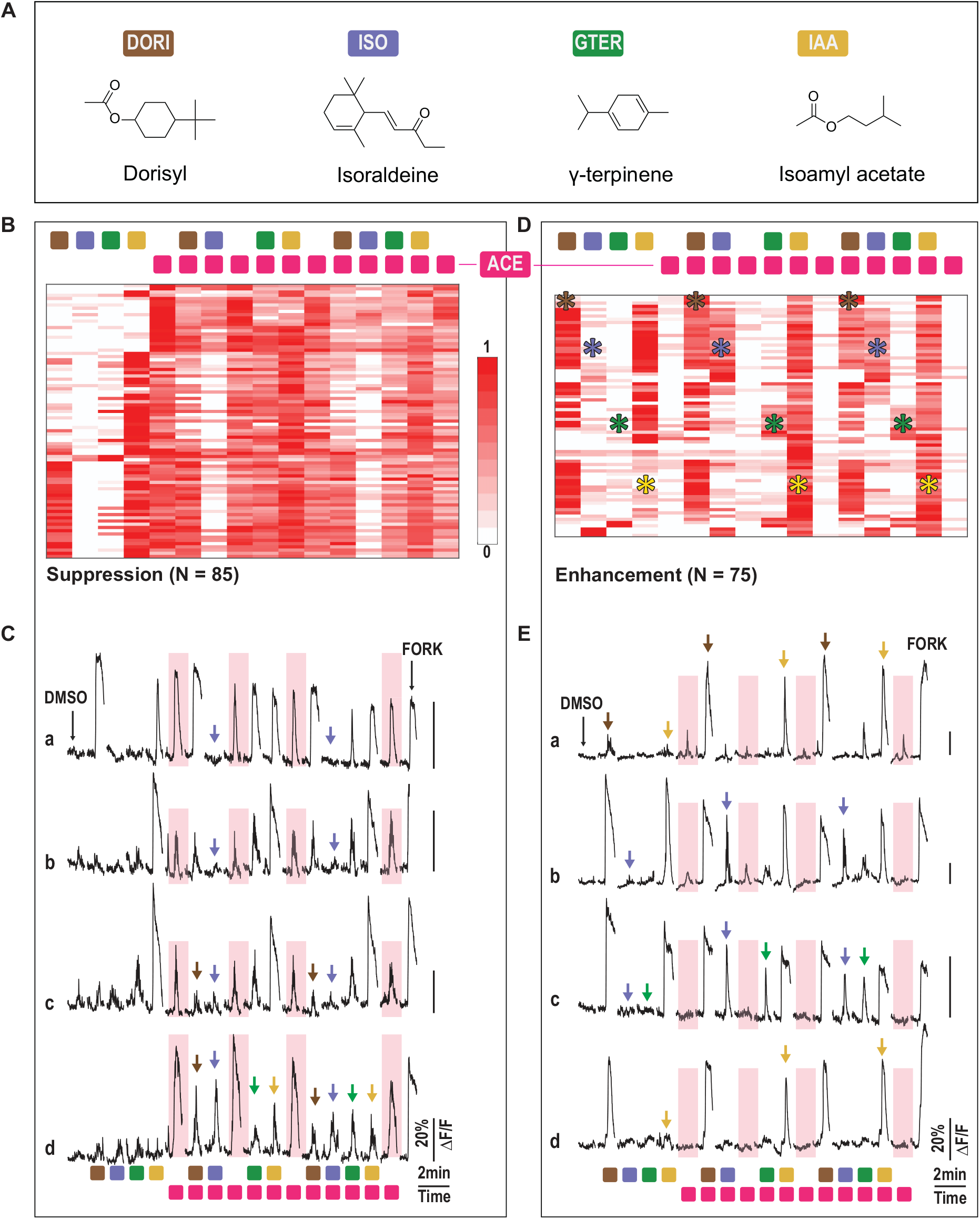
Acetophenone responses can be modulated by multiple odorants. **A.** Chemical structures of odorants tested as modulators. These odorants were chosen to reflect a diversity of chemical properties. **B.** Normalized response heatmap of suppressed OSNs (N = 85). OSNs were first stimulated with 100μM Dorisyl Isoraldeine, γ-terpinene and Isoamyl acetate individually. Each of the odors was then tested against 30μM Acetophenone. Although the level of suppression varied, all OSNs shown here had >30% suppression. Odor stimuli were denoted by colored squares at the top (color coding as in Fig.6A). **C**. Representative time courses of four OSNs showing suppression. Responses to 30μM Acetophenone alone are highlighted by pink rectangles; inverted triangles with different colors indicate suppression/inhibition by corresponding odorants. **D**. Normalized response heatmap of enhanced OSNs (N = 75). The same protocol as for suppression was used here, and again all the OSNs shown here demonstrated >30% enhancement. Asterisks with different colors indicate enhancement by the corresponding odorants. **E**. Representative time courses of four enhanced OSNs. Inverted triangles with different colors indicate enhancement of corresponding odorants.

We also observed evidence for enhancement among these compounds. For example, cell c shown in Fig. 6E did not respond to ISO, gTER or ACE, but both binary odor pairs (ISO/ACE and gTER/ACE) did activate the neuron. A detailed response profile of enhancement is shown in Fig. 6D. Responses before and after are marked with asterisks for comparison. It thus appears that numerous molecules, both those with and without an apparent smell, could act as antagonists at numerous receptors.

## DISCUSSION

Odor suppression in particular has been well documented in psychophysical tests^10–14^, and is a common tool of perfumers. For example, Isoraldeine (also known as γ-methyl ionone) has been known to perfumers and fragrance producers as a masking agent since at least 2001^34^. However, the neural mechanisms underlying these effects have remained obscure. Even the locus of the effect – peripheral, central or both – is undetermined. Although studies using monomolecular odors as stimuli can reveal ligand-receptor relations and categories of odor sensitivity, such studies cannot reveal the mechanisms at play when smelling blends or mixtures of odors, some of which may be highly complex collections of tens to hundreds of compounds. By screening widespread cell-specific responses to more realistic blends of odors we demonstrate here that the receptors themselves are engaged in a variety of modulatory responses, including antagonism, partial agonism and enhancement, before any further processing of the stimulus at higher system levels.

Inhibition or antagonism has been previously observed for single receptor neurons tested with monomolecular compounds^8,18–23^. Moreover, multiple mathematical and biophysical models have been proposed to describe the consequences of mixing two odors that included suppression and/or inhibition^21,35–37^. However, the low throughput of classical methods to assess cell-specific responses to multiple odors has made it very difficult to quantify the incidence of inhibition in practice and therefore tis relevance to perception. Making use of a new high-throughput, high-speed 3D imaging technology (SCAPE microscopy) enabled us to overcome this throughput challenge - surpass this limitation by an order of magnitude - and investigate responses of many neurons in an intact epithelium simultaneously, stimulated with multi component odor blends. These recordings revealed a complex interaction of mixtures of odors at the peripheral sensory level – at odds with the commonly accepted idea of a simple combinatorial encoding of odors at this level. In a strict combinatorial model the responses of a mixture should be approximately a linear summation of the responses to the individual components.

Our results however demonstrate that some odors act as agonists at one receptor and antagonists or partial agonists at others, painting a much more complex picture of how odor sensing leads to perception of mixed odor blends. Additionally an approximately equal number of odors acted as enhancers of other odor responses. Under the simplified conditions of using only a 3 odor blend we nonetheless found evidence of robust modulatory effects between the compounds. Within any one subgroup we observed as much as 38% of the responses being modulated by different components of the mixture, and overall we estimate that a minimum of 22% of responses were modulated, out of 10,000 cells with only 3 odor component blends. Although this is problematic for a simple combinatorial coding strategy it is consistent with recent data from several laboratories that have reported no discernable patterning or topographic arrangement of inputs from olfactory bulb to piriform cortex ^38–40^.

We propose an alternative to a combinatorial code utilizing the modulatory effects observed here (Fig. 7). Without the modulatory effects, the number of receptor patterns could easily be saturated even in the large olfactory receptor family due to the orders of magnitudes higher number of potential ligands (see left panel, model 1). Making conservative estimates that any given odor molecule can activate 3-5 receptors at a medium level of concentration, then a blend of just 10 odors could occupy as many as 50 receptors, more than 10% of the family of human receptors^4,41^. The situation is worsened if, as seems likely, some of these receptors have overlapping sensitivities. This will result in fewer differences between two blends of 10 similar compounds. The number of available unoccupied receptors is further reduced with the addition of each new component, eventually saturating the system and making it impossible to discriminate between complex blends. We note that coffee has over 850 volatiles and Bordeaux wines are reported to have as many as 1100 ^42,43^. However, if each of the ligands (odors) in a blend also inhibits some receptors activated by other components then this removes some of the overlapping receptors, producing increased sparsity (see right panel, model 2). At the same time enhancement introduces receptors insensitive to individual components but activated by the presence of multiple odors, broadening the ligand-binding spectrum of the receptors. Together, these modulatory actions produce additional possible patterns for complex blends to occupy. Therefore we suggest pattern recognition as an alternative to either combinatorial or topographical coding strategies.

**Figure 7.**
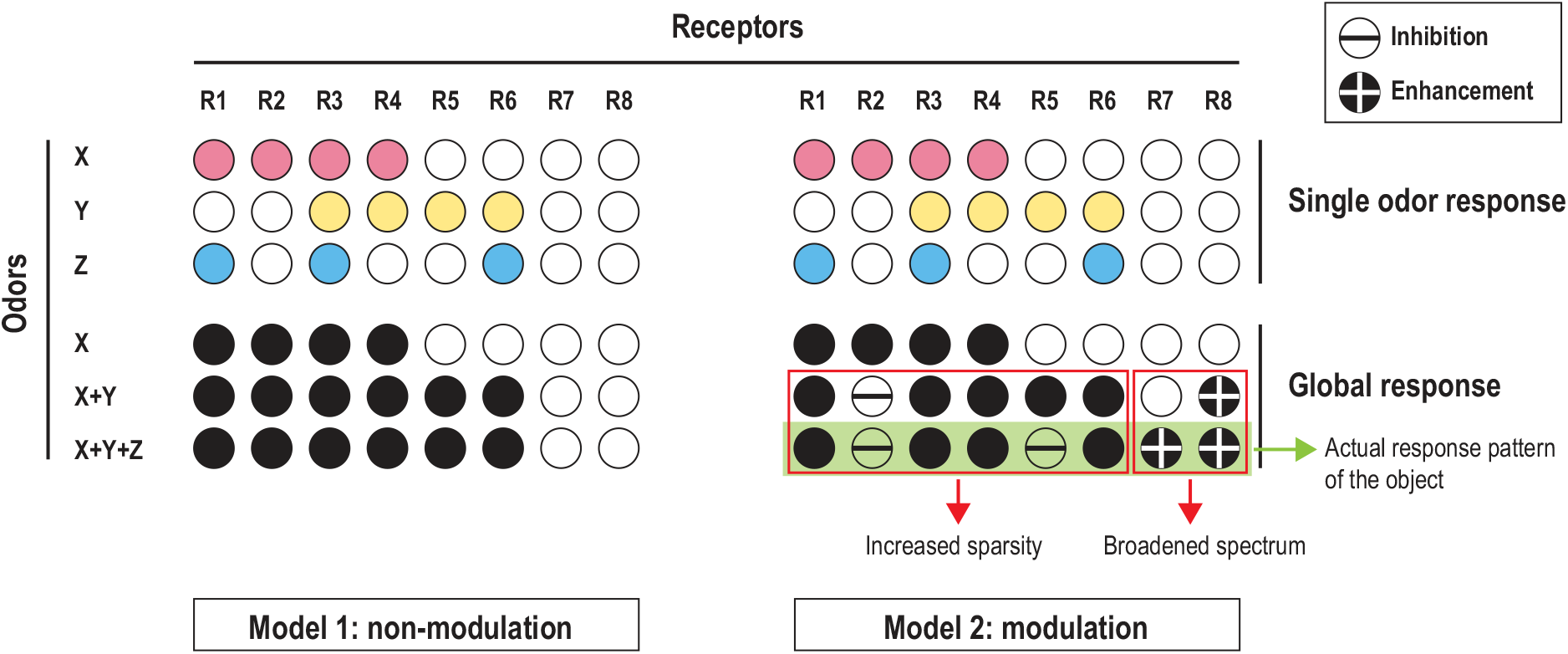
Increased coding capacity through modulation. Two conceptual models are shown to contrast their robustness in odor mixture coding. **Non-modulation model (left):** If we suppose odors X, Y and Z (all monomolecular compounds) can each activate a subset of odor receptors, with odor 1 having a slightly broader spectrum. In this model, mixing odor Y with odor X would recruit two more receptors but adding Z will not produce a different perception, because odor Z recruits no new receptors. On the other hand, the four receptors that have been activated by odor X are ‘locked’ in an activation state, and are not available for detecting component changes in this mixture. **Modulation model (right):** In this model, all receptors are subject to modulation in addition to their activation profile. Under one possible circumstance, similar to what we have observed, mixing odors X and Y results in the inhibition of receptor 2 and the enhancement of receptor 8. Adding odor Z into the mixture inhibits receptor 5 and enhances receptor 7. As a result, the sparsity is increased due to inhibition and the spectrum of odor coding is expanded through enhancement. Together these serve to increase the robustness of pattern detection as a mechanism of perception. This model also implies that ‘silent’ receptors might be as important as the activated ones in pattern recognition of an olfactory object.

Olfaction appears unique in these interactions at the level of primary receptors. In other non-chemosensory sensory systems, (vision, hearing, somatosensory, etc.) there is no demonstrated case of a stimulus activating one type of primary receptor and inhibiting another. Whatever interactions occur between stimulus responses are at higher levels of processing, e.g. red-green color opponency in vision. This raises the crucial question as to whether olfactory processing at higher levels resembles that of other sensory systems or utilizes an alternative strategy. Together with the recent work in piriform cortex suggesting a lack of topographical representation^44^, there is abundant motivation to consider alternative coding strategies that also account for the presence of receptor modulation at the first step of olfactory discrimination.

## Acknowledgements

We would like to thank Zita Peterlin (Firmenich SA), Dongjing Zou (Columbia), Christophe Laudamiel (DreamAir LLC) and Stavros Lomvardas (Columbia) for their discussion and advice in experiment design. We thank Kripa B. Patel (Columbia) and Citlali P. Campos (Columbia) for maintenance of the SCAPE imaging system. We thank Cen Zhang (Columbia) for technical support in mouse breeding and genotyping. We thank Liam Paninski (Columbia) and Eftychios A. Pnevmatikakis (Flatiron) for their help in data analysis. We also thank other members of the Firestein lab and the Hillman lab for their support of this work.

## Author Contributions

LX, SF, WL and EMCH designed the experiments. VV, EMCH and WL designed, constructed and maintained the imaging system. LX and WL designed, constructed the experimental setup and performed the experiments. LX, WL and EMCH analyzed the data. SF and LX prepared the manuscript. All authors discussed and contributed to writing the manuscript.

## METHODS

### Animals

Mice were housed and handled in accordance with protocols approved by Columbia University Institutional Animal Care and Use Committee. OMP-*Cre*-driven GCaMP6f strain was generated by crossing OMP-*Cre* strain (JAX006668) with Ai95D (CAG-GCaMP6f, JAX024105). Male 6 to 8-week old mice with a genotype of OMP-*Cre*^+/−^ GCaMP6f^−/−^ were used for SCAPE Imaging.

### Tissue preparation

Mice were overdosed with anesthetics (ketamine 90 mg·kg^−1^; xylazine 10 mg·kg^−1^, i.p.) and decapitated in accordance with IACUC approved procedures. The head was cut open sagittally and the septum was removed to expose the surface of the olfactory turbinates. Only the right half was used for experiments. The tissue was placed in cold modified Ringer’s solution (mM: 113 NaCl, 25 NaHCO_3_, 5 KCl, 2 CaCl_2_, 3 MgCl_2_, 20 HEPES, 20 Glucose, pH 7.4) for 40min before imaging.

For SCAPE imaging, the right half of a mouse head with the olfactory turbinates exposed was mounted in a custom-designed 3D printed glass bottomed perfusion chamber. The perfusion chamber was designed to control the perfusion flow in the nasal cavity with the inlet at the nostril and the outlet at the throat (Fig. 1A, blue trace and arrows). A small amount of light-cured dental composite (Tetric EvoFlow®, Ivoclar Vivadent) was applied to adhere the tissue to the chamber. During experiments, the tissue was continuously perfused with carboxygenated (95% O_2_, 5%CO_2_) modified Ringer’s solution at room temperature, 0.75ml·min^−1^. Depending on individual differences and the particulars of the tissue mounting, there was some variation in the precise region that was imaged. Typically, the field of view covered the ventral half of either turbinate IIb or III ^45^, and some portion of the neighboring turbinates. Every experiment covered a large and overlapping region of the epithelium (Fig. 1A, yellow rectangle). Analyses are on combined data from all regions recorded, as we saw no apparent differences in regional responses.

### Odorants and odor stimuli

All odorants in this study except Benzyl acetate, Dorisyl, Dartanol, and Isoraldeine (all gifts from Firmenich SA) were from Sigma-Aldrich. In odor set 1, Acetophenone, Benzyl acetate and Citral were first diluted in DMSO to make stock solutions, then subsequently diluted in modified Ringer to 100μM. For two and three-component mixtures, Acetophenone, Benzyl acetate and Citral were mixed then subsequently diluted so that each component has a final concentration of 100μM, with DMSO concentration at 1‰. In odor set 2, Dorisyl, Dartanol and Isoraldeine were mixed at a volume ratio of 45%: 15%: 40% to reproduce the Woody Accord, an accord that has been widely used in the perfume industry. Final concentrations of the three odorants were 148μM, 48μM and 127μM, respectively, both in single odorant solution and in mixtures; DMSO concentration was at 2‰ for enhanced solubility. All other odorant solutions had a final DMSO concentration at 1‰.

Odorants were applied for 30s using a 1260 Infinity HPLC system (Agilent Technologies, Santa Clara, CA, USA) with 2.5min time intervals between stimuli. The adenylate cyclase activator forskolin (50μM, Sigma-Aldrich) was applied at the end of each experiment to assess the viability of OSNs.

### SCAPE imaging

High-speed volumetric imaging of intact epithelium was performed on a custom Swept Confocally-Aligned Planar Excitation (SCAPE) microscope extended from designs described in Bouchard et al. 2015 and Hillman et al. 2019 ^15–17^. SCAPE is a form of light-sheet microscopy, providing low phototoxicity combined with very high-speed 3D imaging of intact samples through a single, stationary objective lens. Briefly, SCAPE’s high-speed 3D imaging is achieved by illuminating the sample with an oblique light sheet through a 1.0 NA objective lens. Fluorescence signal excited by this sheet (extending in y-z’) is collected by the same objective lens (in this case an Olympus XLΜMPLFLN 20XW 1.0 NA water immersion objective with a 2mm working distance). A galvanometer mirror in the system is positioned to both cause the oblique light sheet to scan from side to side across the sample (in the x direction) but also to descan returning fluorescence light. This optical path results in an intermediate, de-scanned oblique image plane which is stationary yet always co-aligned with the plane in the sample that is being illuminated by the scanning light sheet. Image rotation optics and a fast sCMOS camera (Andor Zyla 4.2 PLUS) are then focused to capture these y-z’ images at over 1000 frames per second as the sheet is scanned in the sample in the x direction. Data is then reformed into a 3D volume by stacking successive y-z’ planes according to the scanning mirror’s x-position. All other system parts including the objective and sample stage are stationary during high speed 3D image acquisition.

In this study, the stationary objective in SCAPE system was configured in an inverted arrangement to image underneath the perfusion chamber. The overall magnification of SCAPE microscopy was configured to be 4.66x. A 488nm laser was used for excitation (<1.4 mW at the sample) with a 500 nm long pass filter for the emission path. The system’s sCMOS camera was used at various frame rate for different specific ROI (800-1300 fps). The x-direction scanning step was set to be around 2-3μm to achieve 2-5 volumes per second imaging over a field of view as large as 1000μm × 800μm × 220μm (y-x-z, 1.39×(2-3)×1.1μm per pixel). Each trial was acquired for 75 seconds, and each mouse can be imaged for more than 20 trials with 2.5 min of inter-trial interval.

### Volumetric image data processing

Sample drifts were corrected using custom Matlab code based on NoRMCorre^46^. Since the tissue drifts and motion artifact within each trial is negligible, a single volume from each trial was taken and registered to the template with manual correction when necessary. The same transform matrix was then applied to all the volumes in each trial. After registration, 3D volumetric data from each trial were then divided into multiple 7.7μm depth sub-stacks with 3.3μm spacing to minimize overlapping of neurons. Trials from the same experiment were concatenated and projected into 2D stacks with sum intensity in depth. Constrained Nonnegative Matrix Factorization (CNMF) was applied on each 2D stack to extract single cell calcium activity ^28^.

Neurons with significant tissue drift/deformation were rejected from further analysis. CNMF extracted neuronal time-courses and spatial loci were initially screened with convolutional neural network to exclude neurons with spontaneous activity, motion induced baseline fluctuation or inconsistent responses to repetitive odor stimuli. All the cells were later validated manually to ensure data fidelity. Time courses of each neuron’s GCaMP activity were shown as *ΔF*/*F*, where *ΔF* is the real-time fluorescent intensity change and *F* is the baseline.

### Plotting normalized response heatmaps

Peak responses were normalized to the maximum odorant response of each neuron. Since odorants were administered in a pseudo-stochastic manner, data from different animals were re-aligned before combination. Neurons were then clustered into subgroups based on k-means clustering and sorted according to response intensity and modulation index within each subgroup.

In order to calculate the modulation index (*I_mod_*) in Fig. 3D and 4C, responses to the mixture were linearly corrected based on responses to repeated odor stimuli. The modulation index (*I_mod_*) was calculated as *(d_1_−d_2_)/d_2_*, where *d_1_* is the magnitude of response to the dominant odorant, and *d_2_* is the corrected magnitude of response to the mixture (Fig. 3C). In a control experiment, the variance of *I_mod_* was measured to be 9% (N = 944, 2 mice; see Extended Data Fig. 2), which is comparable to that of conventional calcium imaging data^9^.

## DATA AND SOFTWARE AVAILABILITY

All custom Matlab scripts will be made available upon request. The data that support the findings of this study are available from the corresponding authors upon reasonable request.

## Supplemental Information

**Extended Data Figure 1.**
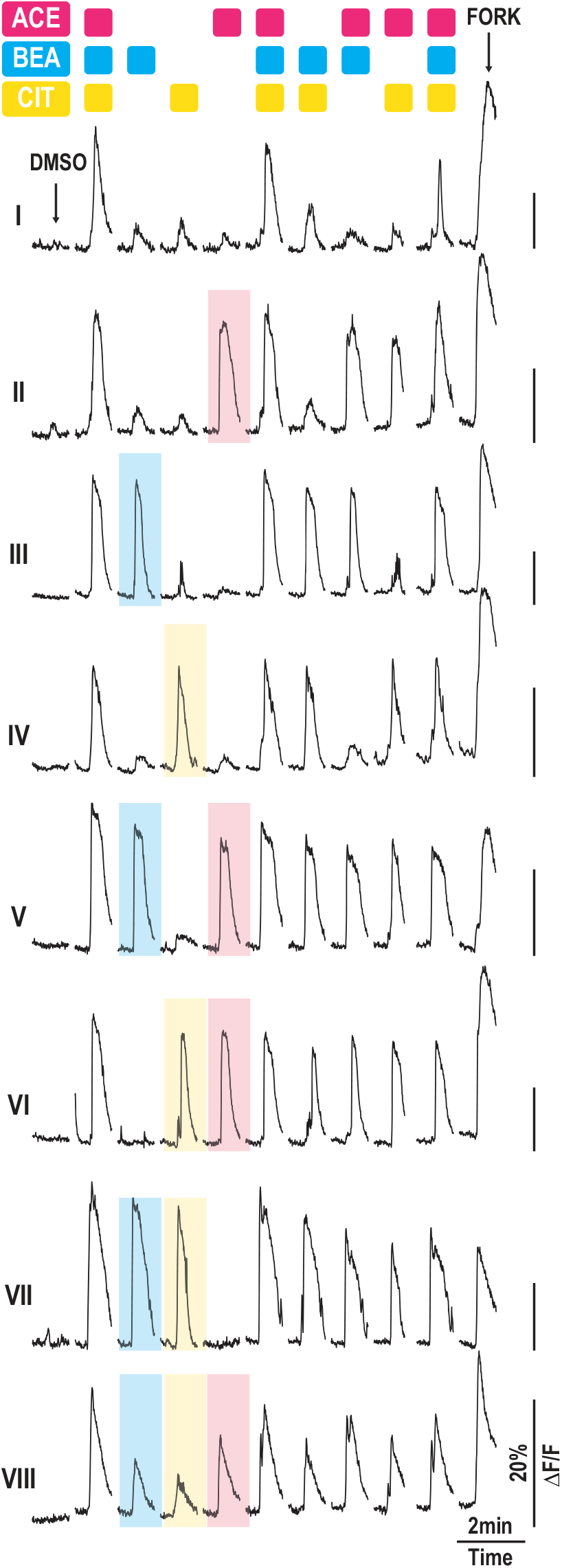
Representative time courses of subgroup I-VIII. A representative OSN was selected from each subgroup to illustrate different response patterns. Responses to 100μM Acetophenone/Benzyl acetate/Citral were highlighted with pink/blue/yellow rectangles, respectively.

**Extended Data Figure 2.**
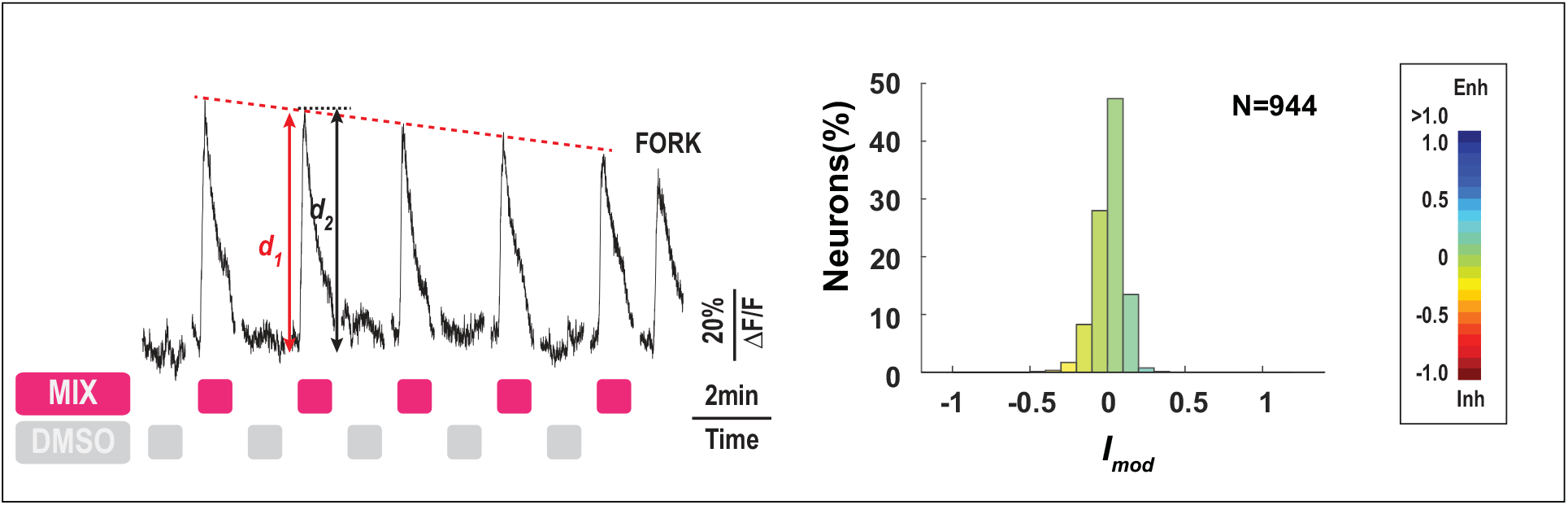
Stable responses to repetitive odor stimuli. A. Representative time course of an individual OSN. Neurons were stimulated repetitively by the ABC mixture (Acetophenone + Benzyl acetate + Citral, 100μM each) and DMSO. Cell viability was verified by 50μM forskolin. B. Histograms showing natural variation of OSN responses. Modulation indices (*I_mod_*) were calculated and plotted using the same method shown in Fig.4B. The results indicate that although there are a few outliers, the actual responses of most OSNs were close to estimation, with a standard deviation of 9% (N = 944, 2 mice).

**Extended Data Figure 3.**
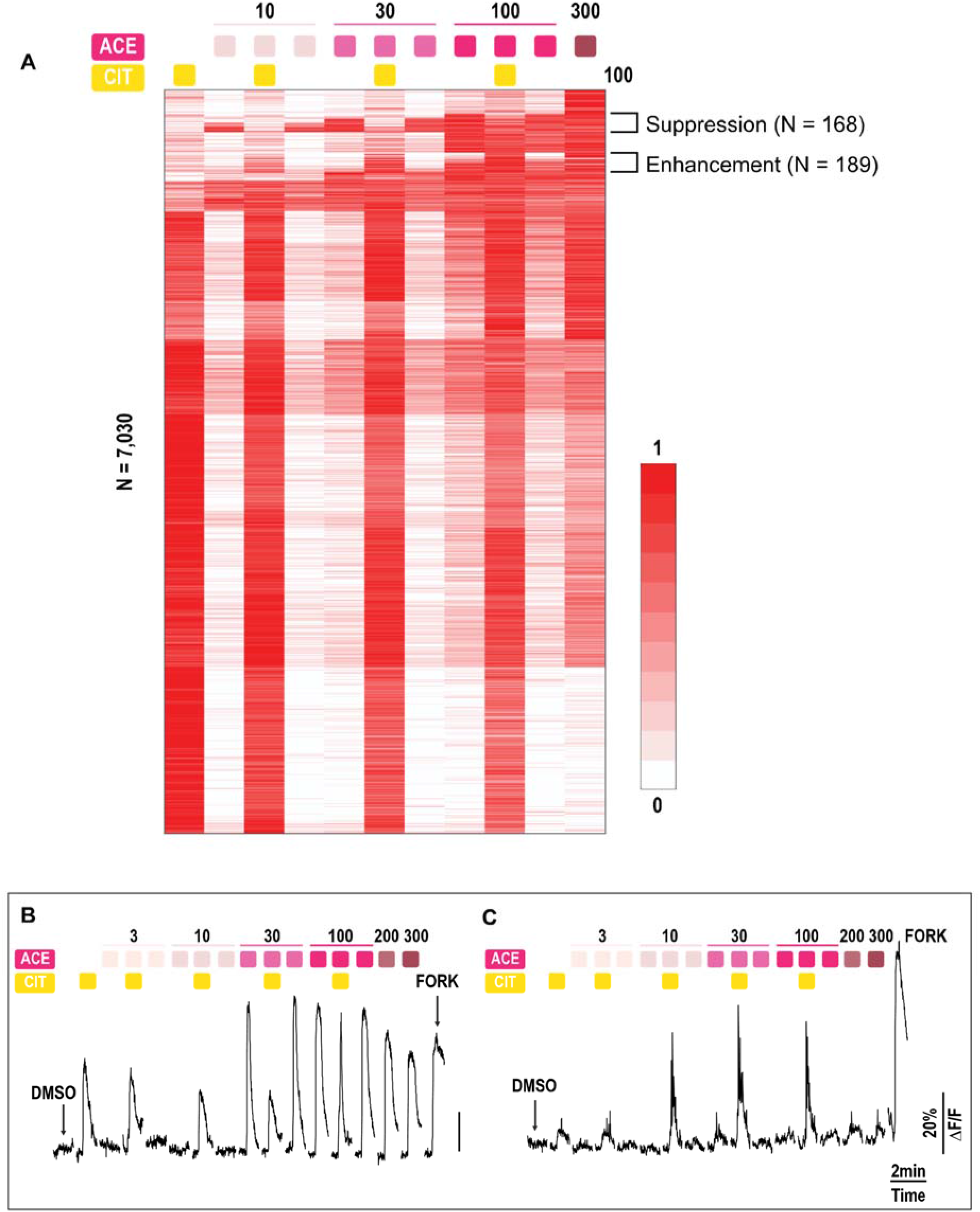
Dose-dependent responses to Acetophenone and Citral. A. Heatmap of normalized peak responses (N = 7030, 3 mice) to Acetophenone and Citral. OSNs were stimulated with increasing concentrations of Acetophenone (10-300μM) in the presence or absence of 100μM Citral. OSNs were sorted based on kmeans clustering results and the modulation index. Brackets of suppression and enhancement indicate neuron subgroups shown in Figure 6A and 6D, respectively. B. Representative time course of an individual OSN showing partial agoinsm by 100μM Citral. C. Representative time course of an individual OSN showing enhancement by 100μM Citral, possibly due to positive allosteric modulation.

**Extended Data Figure 4.**
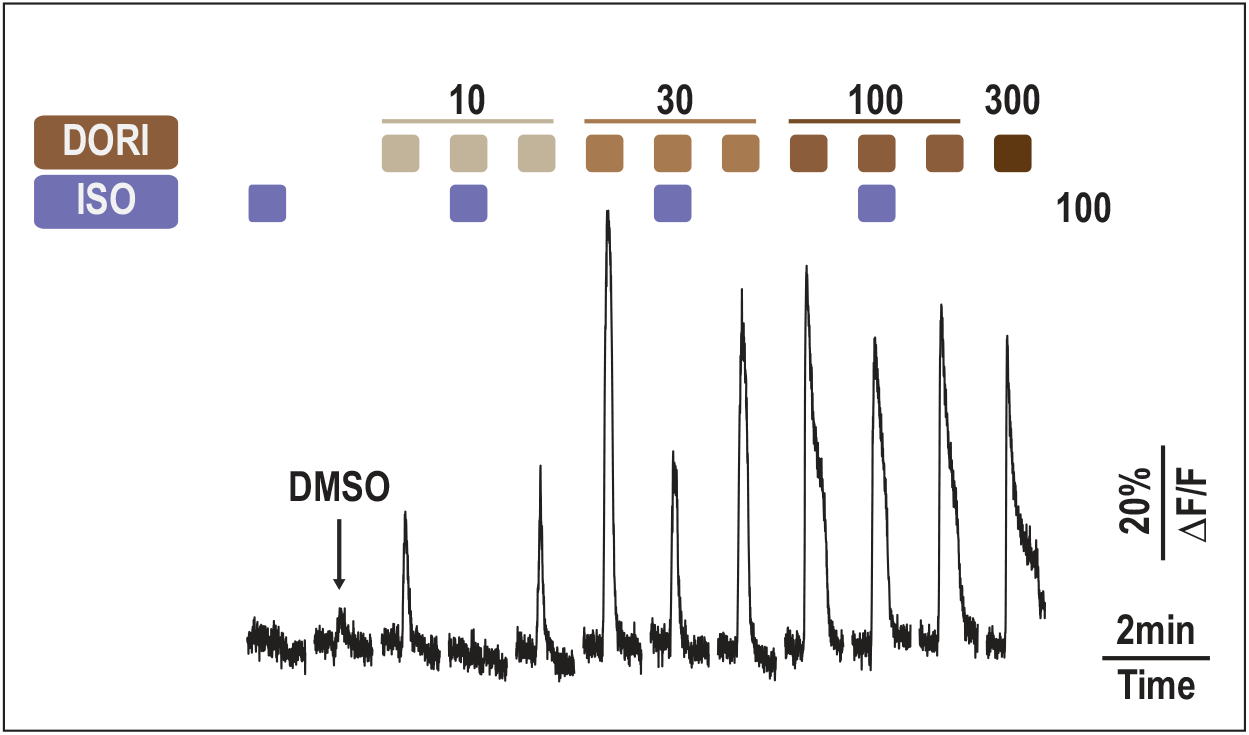
Representative time course of an individual OSN. Neurons were stimulated with increasing concentrations of Dorisyl with or without 100μM Isoraldeine.

**Extended Data Figure 5.**
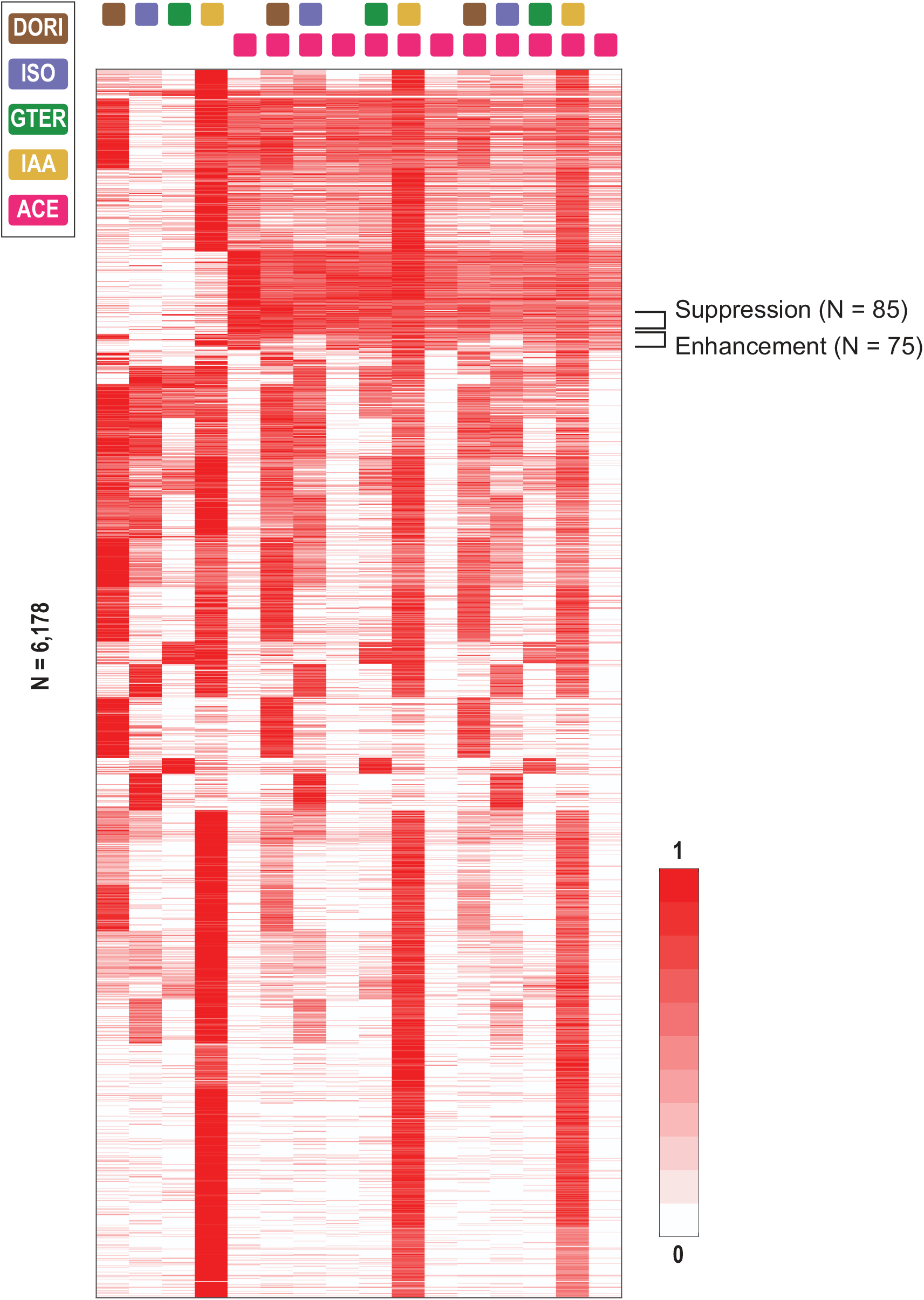
Heatmap of normalized peak responses (N = 6178, 3 mice) to Acetophenone and four candidate modulators. OSNs were first stimulated with 100μM Dorisyl, Isoraldeine, γ-terpinene and Isoamyl acetate without Acetophenone and then with 30μM Acetophenone. Odor stimuli were presented in a pseudo-stochastic manner and re-aligned. OSNs were sorted based on k-means clustering results and sorting of modulation index. Brackets of suppression and enhancement indicate neuron subgroups shown in Figure 7B and 7D, respectively.

**Extended Data Movie 1.**
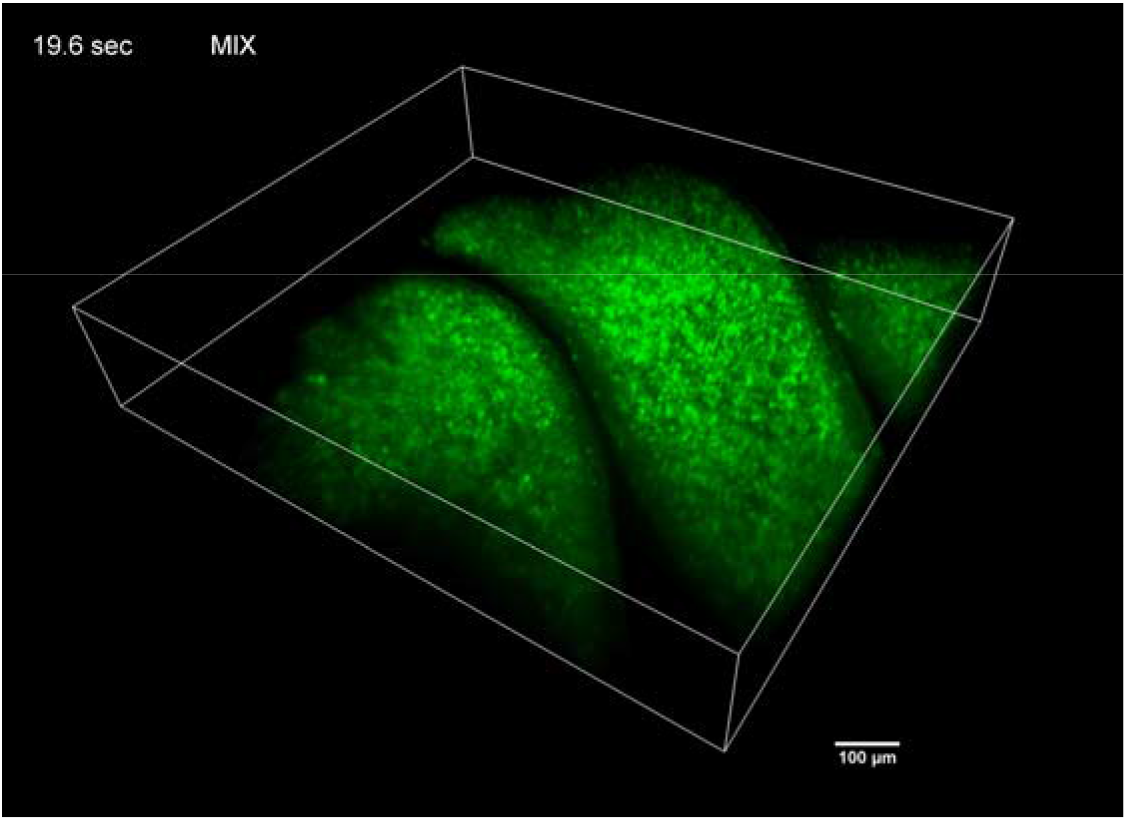
3D rendering video of SCAPE recording olfactory epithelium neuronal GCaMP fluorescence (Turbinate IIa-III) responding to a mixture of Acetophenone, Benzyl acetate and Citral (100μM each). The video playback speed is 5x real time.

## References

1 Buck, L. & Axel, R. A Novel Multigene Family May Encode Odorant Receptors - a Molecular-Basis for Odor Recognition. Cell 65, 175–187, doi:Doi 10.1016/0092-8674(91)90418-X (1991).

2 Zhang, X. M. & Firestein, S. The olfactory receptor gene superfamily of the mouse. Nature Neuroscience 5, 124–133, doi:10.1038/nn800 (2002).

3 Godfrey, P. A., Malnic, B. & Buck, L. B. The mouse olfactory receptor gene family. P Natl Acad Sci USA 101, 2156–2161, doi:10.1073/pnas.0308051100 (2004).

4 Malnic, B., Godfrey, P. A. & Buck, L. B. The human olfactory receptor gene family. P Natl Acad Sci USA 101, 2584–2589, doi:10.1073/pnas.0307882100 (2004).

5 Niimura, Y., Matsui, A. & Touhara, K. Extreme expansion of the olfactory receptor gene repertoire in African elephants and evolutionary dynamics of orthologous gene groups in 13 placental mammals. Genome Res 24, 1485–1496, doi:10.1101/gr.169532.113 (2014).

6 Zhao, H. Functional Expression of a Mammalian Odorant Receptor. Science 279, 237–242, doi:10.1126/science.279.5348.237 (1998).

7 Araneda, R. C., Kini, A. D. & Firestein, S. The molecular receptive range of an odorant receptor. Nature Neuroscience 3, 1248–1255 (2000).

8 Araneda, R. C., Peterlin, Z., Zhang, X., Chesler, A. & Firestein, S. A pharmacological profile of the aldehyde receptor repertoire in rat olfactory epithelium. J Physiol-London 555, 743–756, doi:10.1113/jphysiol.2003.058040 (2004).

9 Peterlin, Z. et al. The Importance of Odorant Conformation to the Binding and Activation of a Representative Olfactory Receptor. Chemistry & Biology 15, 1317–1327, doi:10.1016/j.chembiol.2008.10.014 (2008).

10 Cain, W. S. Odor Intensity - Mixtures and Masking. B Psychonomic Soc 4, 244–244 (1974).

11 Gillan, D. J. Taste-Taste, Odor-Odor, and Taste-Odor Mixtures - Greater Suppression within Than between Modalities. Percept Psychophys 33, 183–185, doi:Doi 10.3758/Bf03202837 (1983).

12 Laing, D. G., Panhuber, H., Willcox, M. E. & Pittman, E. A. Quality and Intensity of Binary Odor Mixtures. Physiol Behav 33, 309–319, doi:Doi 10.1016/0031-9384(84)90118-5 (1984).

13 Kay, L. M., Crk, T. & Thorngate, J. A redefinition of odor mixture quality. Behav Neurosci 119, 726–733, doi:10.1037/0735-7044.119.3.726 (2005).

14 Cashion, L., Livermore, A. & Hummel, T. Odour suppression in binary mixtures. Biol Psychol 73, 288–297, doi:10.1016/j.biopsycho.2006.05.002 (2006).

15 Bouchard, M. B. et al. Swept confocally-aligned planar excitation (SCAPE) microscopy for high-speed volumetric imaging of behaving organisms. Nat Photonics 9, 113–119, doi:10.1038/Nphoton.2014.323 (2015).

16 Hillman, E. M.C., Voleti, V., Li, W. & Yu, H. Light-Sheet Microscopy in Neuroscience. Annu Rev Neurosci 42, 295–313, doi:10.1146/annurev-neuro-070918-050357 (2019).

17 Voleti V, P. K., Li W, Perez-Campos C, Bharadwaj S, Yu H, Ford C, Casper MJ, Yan RW, Liang W, Wen C, Kimura KD, Targoff KL, Hillman EMC. Real-time volumetric microscopy of in-vivo dynamics and large-scale samples with SCAPE 2.0. Nature methods (2019); in press.

18 Firestein, S. & Shepherd, G. M. Neurotransmitter Antagonists Block Some Odor Responses in Olfactory Receptor Neurons. Neuroreport 3, 661–664, doi:Doi 10.1097/00001756-199208000-00001 (1992).

19 Kurahashi, T., Lowe, G. & Gold, G. H. Suppression of odorant responses by odorants in olfactory receptor cells. Science 265, 118–120, doi:10.1126/science.8016645 (1994).

20 Oka, Y., Omura, M., Kataoka, H. & Touhara, K. Olfactory receptor antagonism between odorants. Embo J 23, 120–126, doi:10.1038/sj.emboj.7600032 (2004).

21 Rospars, J. P., Lansky, P., Chaput, M. & Duchamp-Viret, P. Competitive and noncompetitive odorant interactions in the early neural coding of odorant mixtures. J Neurosci 28, 2659–2666, doi:10.1523/JNEUROSCI.4670-07.2008 (2008).

22 Takeuchi, H., Ishida, H., Hikichi, S. & Kurahashi, T. Mechanism of olfactory masking in the sensory cilia. J Gen Physiol 133, 583–601, doi:10.1085/jgp.200810085 (2009).

23 Chaput, M. A. et al. Interactions of odorants with olfactory receptors and receptor neurons match the perceptual dynamics observed for woody and fruity odorant mixtures. Eur J Neurosci 35, 584–597, doi:10.1111/j.1460-9568.2011.07976.x (2012).

24 Chess, A., Simon, I., Cedar, H. & Axel, R. Allelic inactivation regulates olfactory receptor gene expression. Cell 78, 823–834 (1994).

25 Ishii, T. et al. Monoallelic expression of the odorant receptor gene and axonal projection of olfactory sensory neurones (vol 6, pg 71, 2001). Genes Cells 6, 573–573 (2001).

26 Clowney, E. J. et al. Nuclear Aggregation of Olfactory Receptor Genes Governs Their Monogenic Expression. Cell 151, 724–737, doi:10.1016/j.cell.2012.09.043 (2012).

27 Lyons, D. B. et al. An epigenetic trap stabilizes singular olfactory receptor expression. Cell 154, 325–336, doi:10.1016/j.cell.2013.06.039 (2013).

28 Pnevmatikakis, E. A. et al. Simultaneous Denoising, Deconvolution, and Demixing of Calcium Imaging Data. Neuron 89, 285–299, doi:10.1016/j.neuron.2015.11.037 (2016).

29 Gregory, K. J., Sexton, P. M. & Christopoulos, A. Allosteric modulation of muscarinic acetylcholine receptors. Curr Neuropharmacol 5, 157–167, doi:10.2174/157015907781695946 (2007).

30 Burford, N. T. et al. Discovery of positive allosteric modulators and silent allosteric modulators of the mu-opioid receptor. Proc Natl Acad Sci U S A 110, 10830–10835, doi:10.1073/pnas.1300393110 (2013).

31 Garcia-Carceles, J. et al. A Positive Allosteric Modulator of the Serotonin 5-HT2C Receptor for Obesity. J Med Chem 60, 9575–9584, doi:10.1021/acs.jmedchem.7b00994 (2017).

32 Ahn, S. et al. Small-Molecule Positive Allosteric Modulators of the beta2-Adrenoceptor Isolated from DNA-Encoded Libraries. Mol Pharmacol 94, 850–861, doi:10.1124/mol.118.111948 (2018).

33 Stanczyk, M. A. et al. The delta-opioid receptor positive allosteric modulator BMS 986187 is a G-protein-biased allosteric agonist. Br J Pharmacol 176, 1649–1663, doi:10.1111/bph.14602 (2019).

34 Peter Robert Foley, H. D.H., Carl-Eric Kaiser A hard-surface cleaning composition comprising an odor masking perfume. (2001).

35 Marasco, A., De Paris, A. & Migliore, M. Predicting the response of olfactory sensory neurons to odor mixtures from single odor response. Sci Rep 6, 24091, doi:10.1038/srep24091 (2016).

36 Reddy, G., Zak, J. D., Vergassola, M. & Murthy, V. N. Antagonism in olfactory receptor neurons and its implications for the perception of odor mixtures. Elife 7, doi:10.7554/eLife.34958 (2018).

37 Singh, V., Murphy, N. R., Balasubramanian, V. & Mainland, J. D. Competitive binding predicts nonlinear responses of olfactory receptors to complex mixtures. Proc Natl Acad Sci U S A 116, 9598–9603, doi:10.1073/pnas.1813230116 (2019).

38 Ghosh, S. et al. Sensory maps in the olfactory cortex defined by long-range viral tracing of single neurons. Nature 472, 217–220, doi:10.1038/nature09945 (2011).

39 Miyamichi, K. et al. Cortical representations of olfactory input by trans-synaptic tracing. Nature 472, 191–196, doi:10.1038/nature09714 (2011).

40 Sosulski, D. L., Bloom, M. L., Cutforth, T., Axel, R. & Datta, S. R. Distinct representations of olfactory information in different cortical centres. Nature 472, 213–216, doi:10.1038/nature09868 (2011).

41 Young, J. M. et al. Different evolutionary processes shaped the mouse and human olfactory receptor gene families. Hum Mol Genet 11, 535–546, doi:10.1093/hmg/11.5.535 (2002).

42 Flament, I. Coffee flavor chemistry. (John Wiley & Sons, 2001).

43 Robinson, J. & Harding, J. The Oxford companion to wine. (American Chemical Society, 2015).

44 Roland, B., Deneux, T., Franks, K. M., Bathellier, B. & Fleischmann, A. Odor identity coding by distributed ensembles of neurons in the mouse olfactory cortex. Elife 6, doi:10.7554/eLife.26337 (2017).

45 Ressler, K. J., Sullivan, S. L. & Buck, L. B. A zonal organization of odorant receptor gene expression in the olfactory epithelium. Cell 73, 597–609 (1993).

46 Pnevmatikakis, E. A. & Giovannucci, A. NoRMCorre: An online algorithm for piecewise rigid motion correction of calcium imaging data. J Neurosci Methods 291, 83–94, doi:10.1016/j.jneumeth.2017.07.031 (2017).

